# Reinforcement Learning for Antibody Sequence Infilling

**DOI:** 10.1101/2025.08.08.669419

**Authors:** Chak Shing Lee, Conor F. Hayes, Denis Vashchenko, Mikel Landajuela

**Author notes:** These authors contributed equally to this work (Chak Shing Lee, Mikel Landajuela).

## Abstract

We introduce a flexible framework for antibody sequence design that combines an infilling language model with reinforcement learning to optimize functional properties. Our approach leverages a pretrained infilling language model to generate specific antibody regions within full sequences, guided by reinforcement learning to improve desired biophysical characteristics. We implement a range of online learning strategies, exploring both vanilla REINFORCE and Proximal Policy Optimization with Kullback-Leibler (KL) regularization, and demonstrate that KL regularization is essential for maintaining a balance between score optimization and sequence plausibility. We also adapt Direct Reward Optimization to the protein domain by adding a value head to the infilling model, allowing it to learn directly from static (prompt, response, feedback) datasets using a mean-squared error objective. This formulation is particularly useful when only single-trajectory data is available, which is commonly the case for historically collected experimental assays. We evaluate both the online and offline methods across multiple antibody design tasks—including binding affinity, immunogenicity, and expression—and show that our framework improves alignment with measured biophysical properties while outperforming likelihood-only baselines. This integrated online/offline approach enables functionally driven antibody design and provides a scalable toolkit for therapeutic sequence engineering. Code and data are available at https://github.com/LLNL/protein_tune_rl.

## 1 Introduction

Antibody design is vital for therapeutic development due to its precise target specificity, yet traditional experimental workflows struggle to traverse the vast combinatorial sequence space efficiently [5, 13]. Recent breakthroughs in deep learning, such as antibody-specific language models [26, 17, 35] and generative models [1, 44, 22], have unlocked the ability to generate novel antibody variants with structural plausibility and functional potential [24, 15]. However, these approaches often lack mechanisms for targeted optimization: (i) they struggle to infill CDR regions of variable length and context, (ii) they generate sequences based solely on likelihood rather than biophysical or functional objectives, and (iii) they depend on expensive reward computations, limiting sample efficiency.

Antibody design challenges include CDR variability driven by length and sequence diversity [23], the need for conditional infilling in CDR loops like CDR-H3 [1, 35, 15], and the requirement to optimize non-likelihood objectives such as binding affinity or developability under limited computational budgets [19, 40]. Efficient exploration of the sequence space in this context also demands methods that can learn from historical high-quality antibody examples and experimental data without active reward querying.

This paper presents a unified framework combining variable-length antibody infilling and reinforcement learning (RL) for functional optimization:

1. We adopt a transformer-based infilling model, IgLM [35], which enables conditional generation of CDR segments of variable lengths.
2. We apply online RL via Proximal Policy Optimization (PPO) [33] with KL regularization to fine-tune IgLM for downstream reward objectives while maintaining sequence plausibility.
3. We adopt an offline Direct Reward Optimization (DRO) [30] method to align the base IgLM model stably and sample-efficiently with high-quality antibody datasets, and augment it with a value head to enable soft value function learning.
4. We validate our approach on a range of antibody design tasks, including structure-based signatures like beta sheet content and solvent-accessible surface area (SASA), as well as functional objectives such as binding affinity and expression levels, using both historical datasets and simulated environments. Our results demonstrate that the RL-tuned models outperform likelihood-only baselines in both simulated and learned metrics.

The remainder of this paper is organized as follows. Section 2 reviews related work in antibody modeling, infilling, and RL-driven sequence design. Section 3 details the IgLM architecture, infilling mechanism, and online/offline RL training strategies. Section 4 presents experimental evaluations. Finally, Section 5 discusses limitations and avenues for future research.

## 2 Related Work

### Protein language models for sequence infilling

Transformer-based masked and autoregressive protein language models (PLMs) have been highly effective at capturing sequence context. Masked PLMs like ESM [31] learn from vast sequence datasets to predict masked residues. While powerful as scorers, they are less suited for sampling novel or variable-length infillings, and they lack well-defined likelihoods for generated subsequences. Autoregressive models such as ProtGPT2 [12] and ProGen2 [25] can generate full sequences capable of folding into plausible structures. However, these are generally limited to left-to-right generation and cannot directly infill internal regions.

### Controlled infilling and antibody design

To overcome these limitations, models like RITA extend autoregressive PLMs to support bidirectional context for flexible completion [16]. Specialized infilling models, like IgLM [35], are trained directly on antibody sequences with missing complementarity-determining regions (CDRs) to regenerate functionally plausible loops. Another recent model, ReprogBert, applies reprogramming techniques to PLMs like ProtBERT or BERT for multi-CDR infilling without structural templates, showing promise in antibody loop design [24].

### Non-autoregressive and flow-based generative models

Beyond PLMs, alternative generative frameworks have enabled truly flexible infilling. Diffusion-based models (e.g., discrete sequence diffusion or RFdiffusion for backbones) can operate over arbitrary masked positions, facilitating conditional sequence assembly [42]. Flow-based approaches like ProtBFN, trained on UniProt, support direct infilling by sampling the joint distribution of residue positions, matching or exceeding masked LM performance on zero-shot antibody loop completion [3].

### Reinforcement learning for property-directed sequence design

RL has emerged as an effective approach for optimizing sequence properties. DyNA-PPO [2] applies model-based PPO to iteratively propose and evaluate bio-sequence variants, showing improved sample efficiency for DNA, antimicrobial peptides, and binding tasks. Other methods couple pretrained PLMs with RL: PepPPO [6] optimizes peptide–MHC binding using a predictor as oracle. In antibodies, AB-Gen uses a GPT policy fine-tuned via PPO to generate CDR-H3 sequences targeting multi-property constraints and successfully designed HER2-binding loops in silico [45].

### Advanced RL frameworks and continuous fine-tuning

Recent RL methods apply more advanced frameworks. Structured Q-learning incorporates structural priors into Q-learning for antibody design, outperforming standard RL on SARS-CoV binding tasks [10]. Works like [38] and [39] explore *online fine-tuning* of generative models via RL, balancing reward-driven improvement and sampling diversity. These methods use PLM-based or structural reward functions to guide sequence generation towards desired endpoints such as stability, binding affinity, or structural accuracy, while avoiding mode collapse.

### Summary and gap addressed by our work

In summary, modern protein sequence modeling has progressed from generic scoring PLMs to controlled, context-aware infilling, with reinforcement learning increasingly used to optimize functional properties. Yet, existing approaches rarely combine *variable-length infilling in antibody CDR regions* with both online and *offline RL* using experimental data. This integration remains underexplored. Our work fills this gap directly: we support flexible, variable-length antibody infilling via IgLM, and apply RL both online and offline (leveraging experimental datasets). By unifying generation and RL-based property optimization in a single framework, we uniquely enable scalable antibody design that leverages both in silico and empirical data.

## 3 Method

### 3.1 IgLM Architecture & Training

IgLM, introduced by Shuai et al. [36], is a generative language model for antibody sequences that uses an infilling language modeling (ILM) approach. It adopts a decoder-only Transformer (GPT-style) architecture and was trained on a large corpus of antibody sequences (on the order of 10^8^ sequences from the Observed Antibody Space database [21]). Unlike a standard autoregressive (AR) model, which generates a sequence left-to-right using only preceding tokens as context, IgLM can generate a missing segment within a sequence using both the left (upstream) and right (downstream) context. This infilling capability also distinguishes IgLM from a masked language models (MLM): While MLMs are trained to predict masked tokens given a full context, they can not autoregressively generate novel subsequences. Using a bidirectional context in its training objective, IgLM combines the flexibility of bidirectional reasoning with the generative power of an autoregressive model.

Formally, let *X* = (*x*_1_, *x*_2_, …, *x*_*n*_) denote an antibody amino acid sequence of length *n*, where *x*_*i*_ is the amino acid at position *i*. We consider replacing a contiguous span (typically a CDR loop) of this sequence with new content. Suppose *S* = (*x*_*j*_, *x*_*j*+1_, …, *x*_*j*+*m*−1_) is a span of length *m* starting at position *j* in *X*. We remove this span and substitute a special masking token in its place, resulting in a masked context sequence *X*_\*S*_ = (*x*_1_, *x*_2_, …, *x*_*j*−1_, [MASK], *x*_*j*+*m*_, …, *x*_*n*_). IgLM is trained to model the conditional distribution of the missing span given this surrounding context *P*(*S* | *X*_\*S*_). In the following, we denote the IgLM model with parameters *θ* as *π*_*θ*_(*S* | *X*_\*S*_). For clarity of notation, let *X*_*S*_ = (*x*_1_, *x*_2_, …, *x*_*j*−1_, *S, x*_*j*+*m*_, …, *x*_*n*_) denote the complete sequence formed by inserting the span *S* into its original position within the context.

To learn *π*_*θ*_(*S* | *X*_\*S*_) self-supervised, IgLM uses an autoregressive training strategy on an augmented sequence that includes both the context and the masked-out span. In each training example, we construct a token sequence **X** by concatenating the masked context, a separator token, and the masked span (as the “answer”). Specifically, the augmented sequence is formatted as:

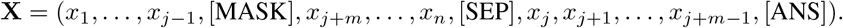

In this formulation, [SEP] is a special separator token that delineates the context portion from the infill segment, and [ANS] is a token marking the end of the inserted span. The model is trained to minimize the standard autoregressive negative log-likelihood over this sequence. That is, for each position *i* in **X**, the model predicts token **X**_*i*_ from all previous tokens **X**_*<i*_, and the training loss is:

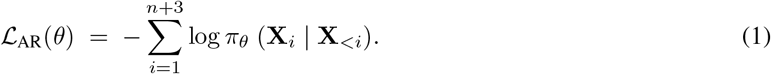

This infilling training scheme allows the model to redesign arbitrary regions of an antibody sequence by generating new amino acid segments that are contextually plausible and diverse.

### 3.2 RL-Based Alignment Framework

We treat antibody sequence generation as an RL-based alignment problem. Let *π*_ref_(*S* | *X*_\*S*_) denote the original pretrained IgLM model (trained via loss (1)), and let *π*_*θ*_(*S* | *X*_\*S*_) be our parameterized policy that we seek to align using a reward function *r*(*X*). We assume access to a distribution *D* of masked contexts *X*_\*S*_ (“prompts”) from which we can sample. We optimize the following KL-regularized expected reward objective:

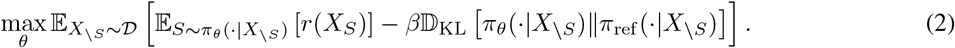

Here *β >* 0 controls the strength of KL regularization, and D_KL_ indicates KL divergence. This objective encourages generating antibody spans that score highly on the reward while staying close to the pretrained model, preserving biological plausibility and preventing unrealistic drift or degenerate sequences. Such regularization has been shown effective in RL fine-tuning of language models, improving training stability and mitigating “reward hacking”, where high proxy reward arises from exploitation rather than true functional improvement [4]. For intuition, consider a simplified scenario where the optimal policy has a closed-form solution. Figure 4 in the appendix illustrates how the temperature parameter *β* shapes the optimal policy’s trade-off between reward maximization and staying near the reference policy.

### 3.3 Online RL Fine-Tuning (Oracle-Driven)

In this section, we assume access to an *online oracle*, a scoring function that can immediately evaluate any complete antibody sequence for quality. In each training step, the model observes a masked context *X*_\*S*_ ∼ *D*, samples a candidate span *S*, and forms a full sequence *X*_*S*_. The oracle then returns a reward *r*(*X*), which is used to fine-tune the model’s policy *π*_*θ*_(·|*X*_\*S*_). The overarching objective is to maximize the expected reward under this real-time feedback loop.

#### REINFORCE (Policy Gradient)

We begin with the REINFORCE algorithm [43], which yields an unbiased Monte Carlo estimator of the gradient of the expected reward:

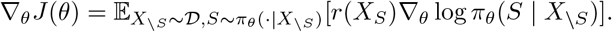

In practice, this expectation is approximated by averaging over a batch of masked contexts and infilling spans. Each update increases the likelihood of spans that received higher rewards. Although REINFORCE is unbiased, it suffers from high variance, sensitivity to learning rate, sample inefficiency, and instability, especially in high-dimensional action spaces or long-horizon tasks. Variance reduction techniques, like subtracting a baseline (e.g. value-function estimate), can reduce noise and speed up learning without biasing the gradient estimate.

#### Proximal Policy Optimization (PPO)

PPO stabilizes policy updates by clipping the surrogate objective to prevent large departures from the previous policy. Specifically, for output span *S* in context *X*_\*S*_, define the probability ratio 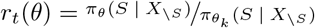, where 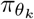 is the fixed policy used to sample the current batch. Following the RLHF framework [37], we also employ a *KL-regularized reward r*^KL^(*X*_*S*_) = *r*(*X*_*S*_) − *β* log *π*^*θ*^ (*S* | *X*^*\S*^)*/π*_ref_ (*S* | *X*_*\S*_). The clipped surrogate PPO objective is:

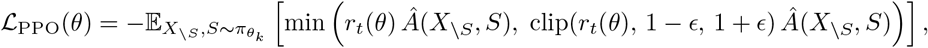

where 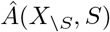 is the advantage estimate (e.g. via GAE). This min-clip structure ensures policy updates remain within [1 − *ϵ*, 1 + *ϵ*], effectively enforcing a first-order trust region that limits policy drift and discourages updates that would push *π*_*θ*_ too far from the reference. The explicit KL term further anchors the policy to the pretrained model, supporting stable RL fine-tuning in RLHF contexts.

### 3.4 Offline RL via Direct Reward Optimization (DRO)

Generating high-reward antibody sequences often requires expensive evaluations (e.g., molecular simulations or expert scores). To avoid repeatedly querying a reward oracle during training, we adopt an *offline* RL approach, leveraging a fixed dataset of 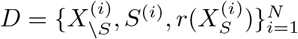 triples. Offline RL enables policy training from historical data without active sampling, which is crucial in protein generation tasks where online reward functions are computationally expensive or infeasible [27]. Recent works, such as DPO for protein optimization [28] and ORPO [18], have demonstrated that preference-or reward-aligned policies can be effectively learned in this offline setting.

#### Deriving the Optimal DRO Policy and Value Function

Starting from the KL -regularized RL objective (2), the optimal policy can be derived in closed form: 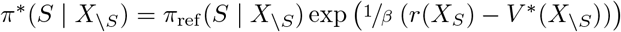, where the *soft value function* is defined by 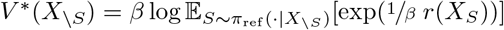 (see Richemond et al. [30]). Figure 4 in the appendix visually illustrates how *π*^∗^ and *V* ^∗^ vary under different values of *β* in a simplified setting. This yields the *optimality condition*:

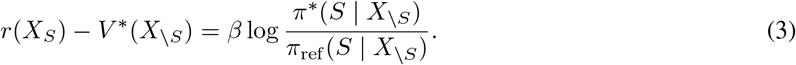

Here, *V* ^∗^(*X*_\*S*_) acts like a soft maximum over reward values under the reference policy. As *β* → 0, *V* ^∗^ approaches max_*S*_ *r*(*X*_*S*_), causing *π*^∗^ to concentrate its mass on the highest-reward spans while remaining anchored to *π*_ref_. This provides a principled, controlled trade-off between exploration and adherence to the pretrained model—ideal for biologically plausible sequence modification.

#### DRO Training Objective

To approximate the analytically derived optimal policy *π*^∗^, we *augment the base IgLM policy network π*_*θ*_ *with an attached value head V*_*φ*_, and train both components by minimizing a mean-squared error loss derived from the optimality condition (3):

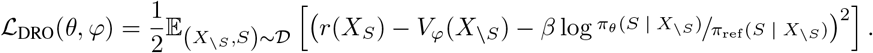

This objective regresses the log-probability ratio of *π*_*θ*_ and *π*_ref_ against the normalized reward, encouraging *π*_*θ*_ to approximate the optimal reward-transformed policy *π*^∗^. This framework, introduced in [30], has been demonstrated as a simple and effective single-trajectory training method that avoids the need for preference-pair data.

Using a minibatch 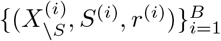, the gradient for the policy *θ* becomes:

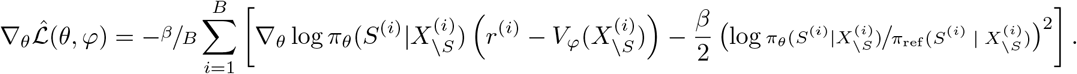

while the update for the value network φ is:

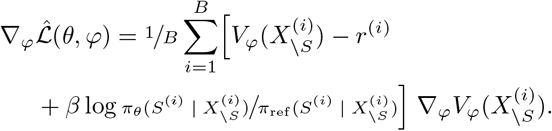

This training procedure mirrors REINFORCE-like policy updates with shaped rewards, while enforcing proximity to the reference distribution. The value network ensures reliable estimation of the soft-normalized reward baseline, making the policy updates stable and consistent [30].

## 4 Experiments

We evaluate our antibody sequence design approach in both online and offline reinforcement learning settings, reflecting the availability of reward feedback. When an interactive reward function is available (e.g., a computational proxy for structural properties), an on-policy RL algorithm can be used to directly optimize sequences in silico. In contrast, real-world antibody design data usually come as fixed historical measurements (a “single-trajectory” dataset of sequences and their properties), making offline RL techniques necessary [40]. Below, we first present results in an idealized online setting with structure-based rewards, and then turn to offline RL results on experimental datasets. Throughout, we use the original IgLM as a baseline to assess the added value of RL fine-tuning and focus on the CDR-H3 loop design. All experiments were executed on two identical compute nodes (a total of 8 GPUs), each housing four NVIDIA Tesla V100 accelerators (Volta architecture, 16GB HBM2 per GPU).

### 4.1 Online RL with Structure-Based Rewards

#### Setup

In the online setting, the agent can query a reward oracle for any generated sequence. We defined three in silico tasks with known structure-based reward functions: (i) Beta sheet percentage - reward proportional to the fraction of residues in *β*-sheet conformation [9], (ii) Solvent Accessible Surface Area (SASA) - reward based on the SASA of the protein’s predicted structure, and (iii) Predicted RMSD (pRMSD) - negative reward equal to the predicted structural deviation of the sequence from a target backbone (lower pRMSD implies the sequence folds closer to a given template). These proxy rewards are computed using the structure prediction tool IgFold [32]. For each task, we fine-tuned the infilling language model using REINFORCE and PPO at different KL regularization strengths *β* (0.01, 0.1, and 1.0) to control exploration-exploitation trade-offs. We also report the performance of our offline RL method DRO when applied in a batch mode using trajectories sampled from the initial policy and labeled with the same structure-based rewards offline (to mimic an offline scenario). The beta sheet percentage and the SASA score tasks have been previously used in [14]. The distribution *D* of masked context is the same as in [14].

#### Results

We report quantitative results for all three structure-based optimization tasks in Tables 1, 2, and 3. Each table shows the average reward scores on a held-out set of validation sequences, along with log-likelihoods from three pretrained language models (IgLM, ProtGPT2, and ProGen) to assess the plausibility and fluency of the generated sequences. Figure 2 (top row) visualizes the distributions of reward scores for PPO-trained models across different KL regularization strengths, while the bottom row shows the corresponding IgLM log-likelihood distributions. Similar plots for DRO are provided in Figure 5 of the Appendix. These plots illustrate the trade-off between structural optimization and adherence to the pretrained policy. Finally, Figure 3 presents predicted 3D structures for sequences generated by different models given the same prompt, offering qualitative insight into how PPO training at varying KL strengths affects folding behavior and structural quality.

**Table 1.**
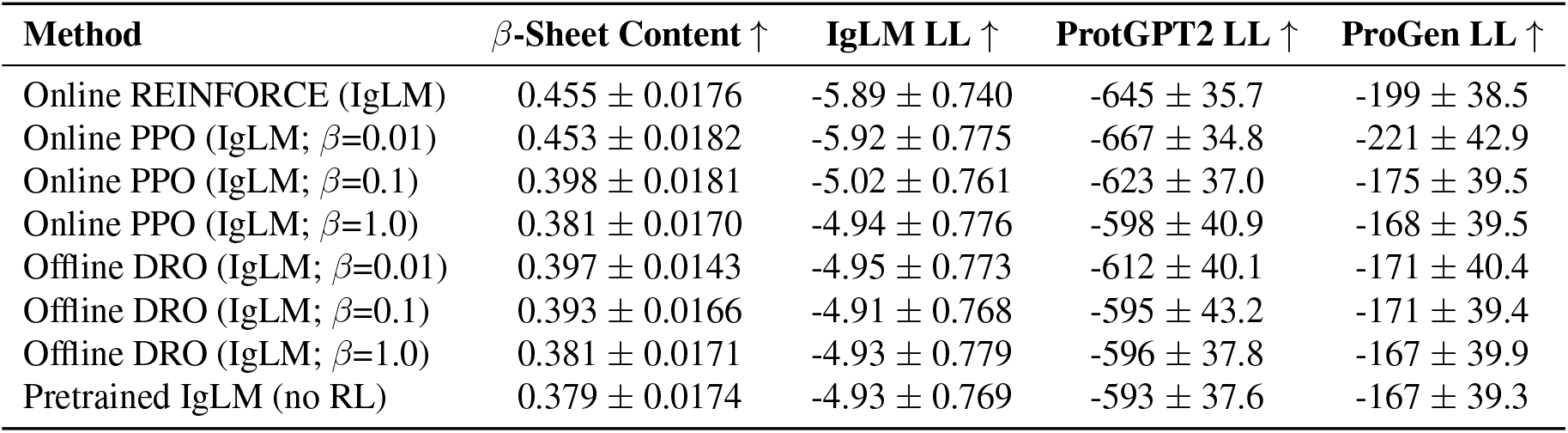
Beta Sheet Optimization: Comparison of methods on average *β*-sheet content in generated sequences. Higher is better. Log-likelihoods under IgLM, ProtGPT2, and ProGen are provided to assess sequence plausibility and fluency.

**Table 2.** SASA Optimization: Comparison of methods on average solvent-accessible surface area (SASA) in folded structure predictions. Higher is better. Language-model log-likelihoods reflect naturalness of sequences under pretrained models.

| Method | SASA ↑ | IgLM LL ↑ | ProtGPT2 LL ↑ | ProGen LL ↑ |
| --- | --- | --- | --- | --- |
| Online REINFORCE (IgLM) | 13310 ± 223 | -5.82 ± 0.704 | -666 ± 33.0 | -207 ± 38.3 |
| Online PPO (IgLM; $\beta=0.01$) | 13320 ± 193 | -5.91 ± 0.728 | -679 ± 34.9 | -209 ± 42.6 |
| Online PPO (IgLM; $\beta=0.1$) | 12764 ± 200 | -5.28 ± 0.733 | -657 ± 30.6 | -182 ± 40.5 |
| Online PPO (IgLM; $\beta=1.0$) | 12504 ± 190 | -4.99 ± 0.760 | -608 ± 37.8 | -169 ± 39.1 |
| Offline DRO (IgLM; $\beta=0.01$) | 12798 ± 211 | -5.31 ± 0.708 | -660 ± 28.8 | -186 ± 39.8 |
| Offline DRO (IgLM; $\beta=0.1$) | 12690 ± 207 | -5.22 ± 0.720 | -649 ± 27.1 | -180 ± 37.8 |
| Offline DRO (IgLM; $\beta=1.0$) | 12486 ± 207 | -4.96 ± 0.772 | -604 ± 37.6 | -168 ± 39.5 |
| Pretrained IgLM (no RL) | 12464 ± 199 | -4.93 ± 0.769 | -593 ± 37.6 | -167 ± 39.3 |

**Table 3.** Predicted RMSD Minimization (pRMSD): Comparison of methods on structural deviation from a target backbone (lower is better). Also shown are log-likelihoods from pretrained LMs (IgLM, ProtGPT2, ProGen) to assess sequence plausibility.

| Method | pRMSD ↓ | IgLM LL ↑ | ProtGPT2 LL ↑ | ProGen LL ↑ |
| --- | --- | --- | --- | --- |
| Online REINFORCE (IgLM) | $0.349 \pm 0.0636$ | $-4.73 \pm 0.809$ | $-535 \pm 39.2$ | $-158 \pm 39.7$ |
| Online PPO (IgLM; $\beta=0.01$) | $0.283 \pm 0.0500$ | $-4.82 \pm 0.820$ | $-544 \pm 46.3$ | $-158 \pm 40.5$ |
| Online PPO (IgLM; $\beta=0.1$) | $0.351 \pm 0.0615$ | $-4.79 \pm 0.813$ | $-542 \pm 42.6$ | $-160 \pm 40.5$ |
| Online PPO (IgLM; $\beta=1.0$) | $0.378 \pm 0.0660$ | $-4.90 \pm 0.780$ | $-584 \pm 34.7$ | $-165 \pm 39.4$ |
| Pretrained IgLM (no RL) | $0.386 \pm 0.0639$ | $-4.93 \pm 0.769$ | $-593 \pm 37.6$ | $-167 \pm 39.3$ |

**Figure 1.**
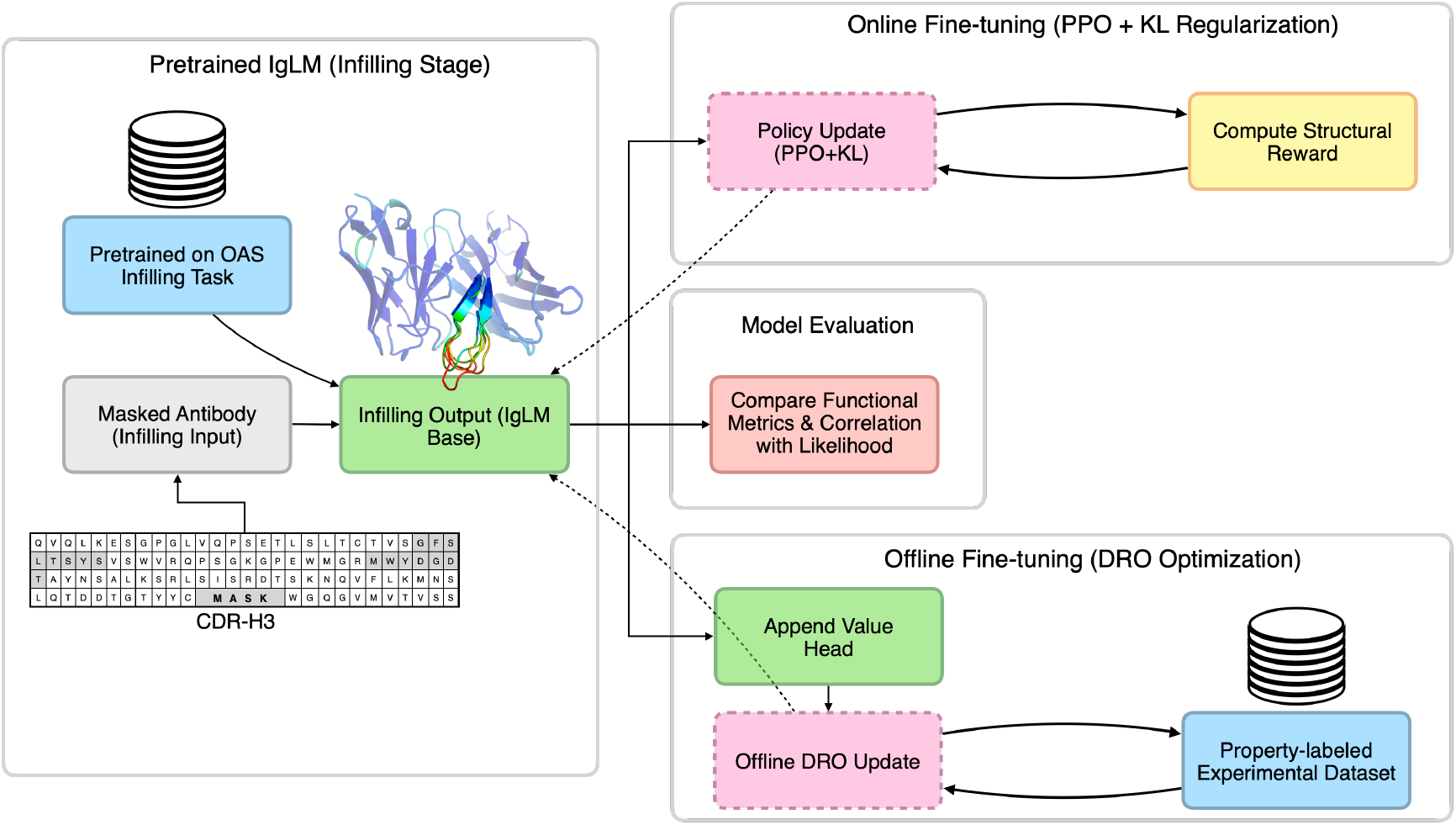
Fine-tuning pipeline overview: pretrained IgLM infills masked CDR-H3 prompts, then fine-tuned using RL via one of two approaches, offline Direct Reward Optimization with labeled binding datasets or online Proximal Policy Optimization incorporating KL-regulation and structure-based rewards. The resulting model is evaluated on functional binding metrics and sequence plausibility.

**Figure 2.**
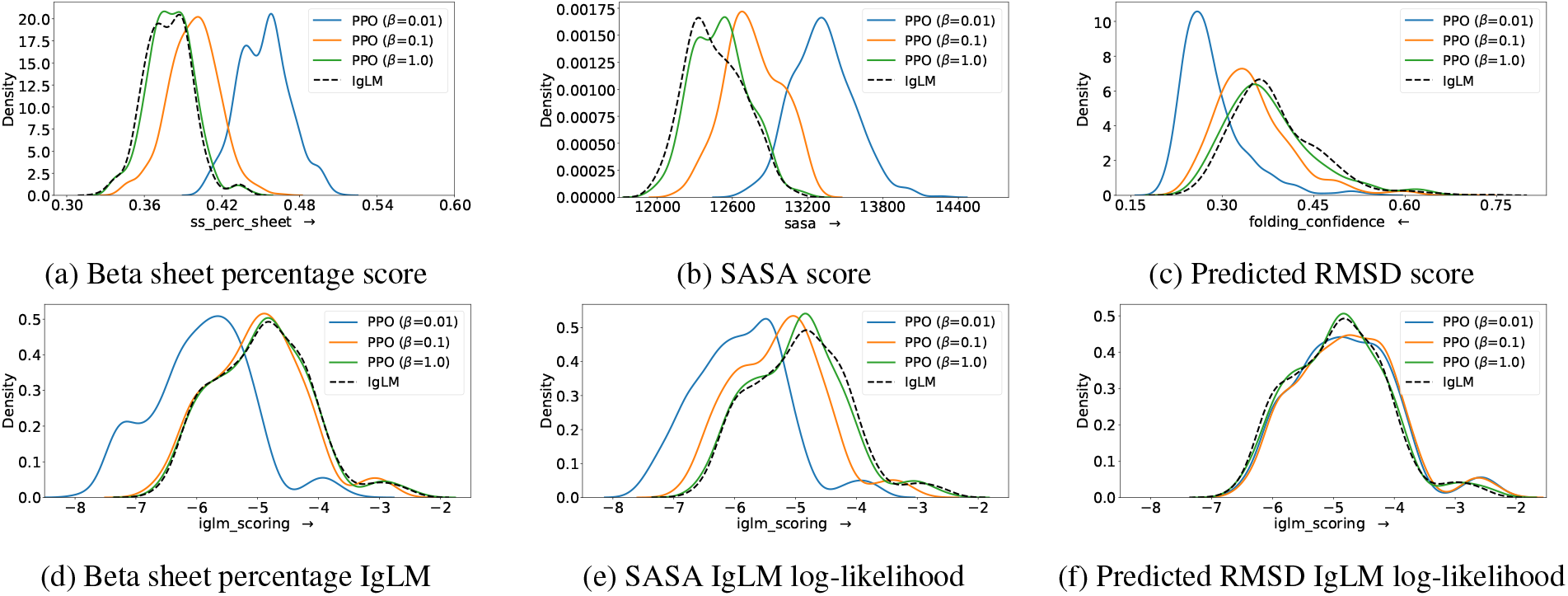
Distribution of scores and IgLM log-likelihoods on the evaluation set for models fine-tuned using PPO on various structure-based reward tasks.

**Figure 3.**
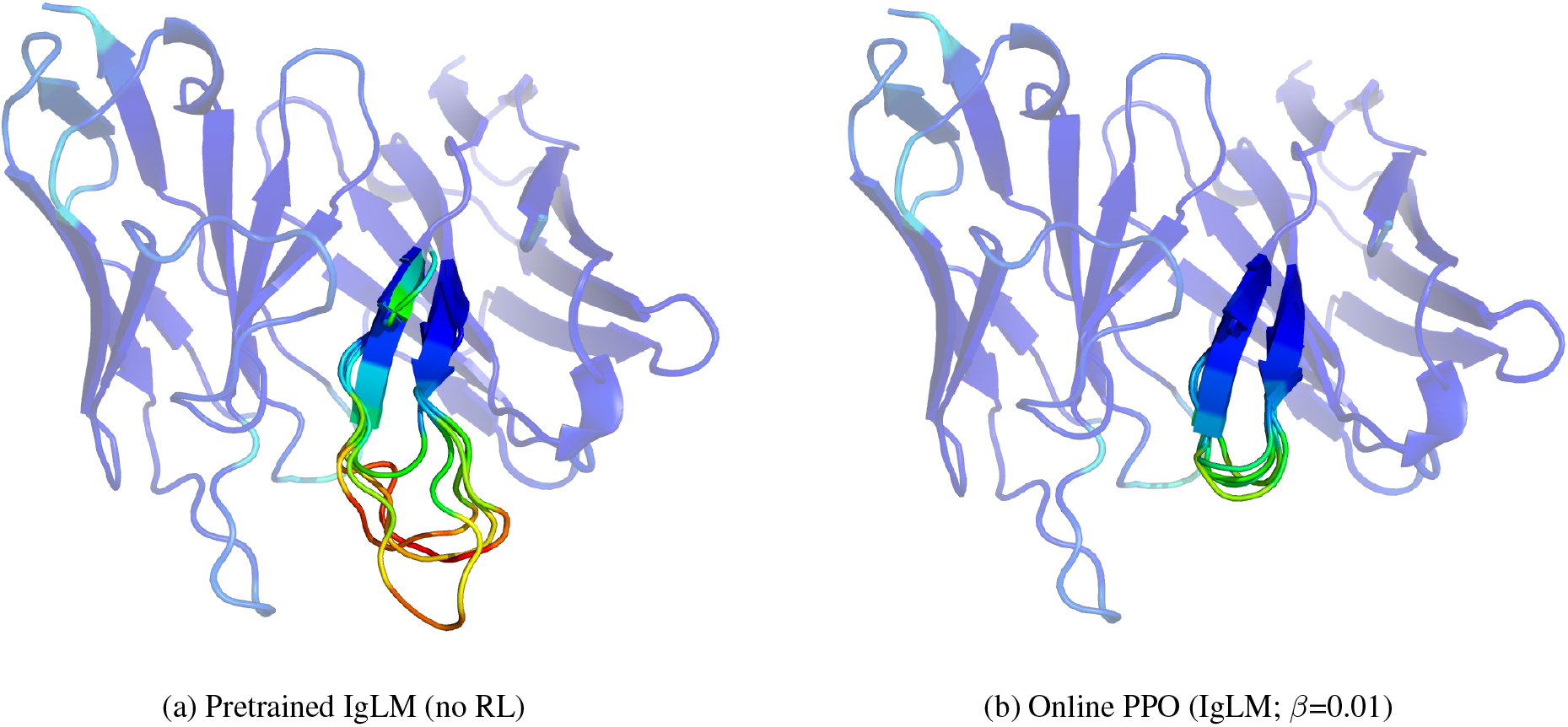
Side-by-side CDR-H3 structural predictions for the same prompt, showing the sequence with lowest pRMSD, one from the pretrained IgLM and one from the PPO fine-tuned model with *β*=0.01. Colder colors indicate lower pRMSD, i.e., closer to the target structure.

Overall, the results support several key findings. First, the KL regularization coefficient *β* plays a critical role in controlling the exploration-exploitation trade-off, as evident from the distribution plots, which mirror the behavior of the theoretical optimal policy in Figure 4 (Appendix). Second, all RL fine-tuned models consistently outperform the vanilla IgLM baseline across all tasks, confirming the efficacy of RL for optimizing antibody sequences. While REINFORCE achieves higher reward scores than PPO and DRO in the beta sheet and SASA tasks, this comes at the cost of generating sequences with lower plausibility, as measured by language model log-likelihoods. Online RL methods (PPO and REINFORCE) generally outperform the offline DRO method, highlighting the value of interacting with the reward oracle during training. Interestingly, in the pRMSD task, all models improve the target reward while maintaining similar IgLM log-likelihoods, suggesting a lack of trade-off in this setting. This alignment between structural accuracy and plausibility may reflect the inductive biases or training data distribution of the IgLM model.

**Figure 4.**
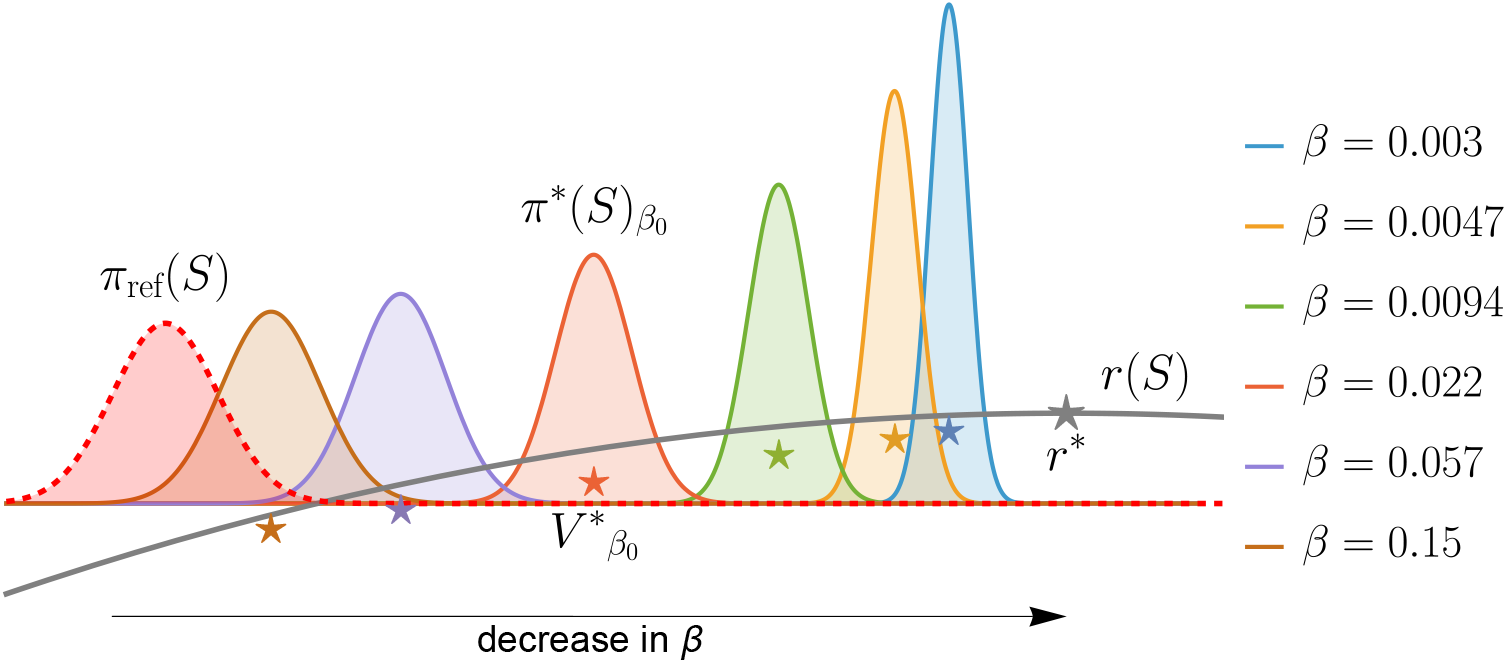
Theoretical optimal policy *π*^∗^(*S*)_*β*_ under different values of reward temperature *β*, assuming a Gaussian reference policy *π*_ref_(*S*) = *N*(*S* | *θ*_ref_ = −0.7, *σ*_ref_ = 0.1) (dashed red line) and a quadratic reward function *r*(*S*) = − (*S* − 1.0)^2^ + 2. The soft value function *V* ^∗^ is presented with a ⋆ symbol, and located at the mean of each optimal policy for each *β*. As *β* decreases, the policy becomes more concentrated around high-reward regions, illustrating the effect of temperature on policy sharpness.

## 4.2 Offline RL on Experimental Datasets

### Setup

In practical antibody design scenarios, wet lab experiments are costly and time-consuming, making online interaction impractical. Therefore, we focus on offline RL methods that rely solely on existing experimental datasets. We curated nine diverse antibody datasets encompassing biologically relevant properties (see Appendix Table 5), including binding affinity (e.g., equilibrium dissociation constant *K*_*D*_ or *IC*_50_ against various antigens), developability or expression measurements (e.g., soluble expression yield), and immunogenicity scores. Each dataset consists of sequences, typically heavy-chain CDR3 variants or full-length antibodies, with corresponding experimental measurements, formalized as 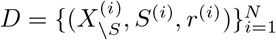 where 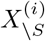 is the sequence context, *S*^(*i*)^ is the CDR-H3 segment, and *r*^(*i*)^ is the measured property (e.g., binding affinity). We divide each dataset into training (80%), validation (10%), and test (10%) splits.

Model performance is evaluated by the Spearman correlation *ρ* between the model’s log-likelihood scores for the sequences and the experimental measurements. A higher correlation implies that the model learns to assign higher probabilities to sequences exhibiting the desired functional property. In practical terms, this means the model can propose new sequences that are expected to perform well in wet lab validation based on past experimental data.

### Results

Table 4 presents the Spearman correlation (*ρ*) between model-assigned log-likelihoods and actual experimental measurements for each antibody dataset (test set). The table compares the pretrained IgLM baseline with the Offline DRO fine-tuned model (*β*=0.01). Across nearly all datasets, Offline DRO significantly improves correlation relative to the pretrained baseline, often transforming near-zero or negative correlation (e.g., −0.09 for MITLL AlphaSeq binding) into moderate-to-strong alignment (e.g., *ρ* = 0.45, *p <* 0.001). Improvements are consistent across expression datasets (e.g., G6 expression: *ρ* rises from −0.01 to 0.41) and immunogenicity (Marks 2021: *ρ* from 0.17 to 0.46, *p <* 0.01). Although IgLM log-likelihoods modestly decrease following DRO fine-tuning, indicating only a slight shift from the base language model, the decline is minimal, and it is paired with significantly improved Spearman correlation. This suggests that fine-tuning enhances prioritization of functional properties while largely preserving sequence plausibility. Overall, these results validate that offline DRO fine-tuning enables the model to leverage labeled experimental data for effective sequence prioritization, outperforming the vanilla pretrained model across diverse datasets and property categories.

**Table 4.** Offline RL Evaluation on Experimental Datasets. Spearman’s *ρ* between model log-likelihood and the experimental measurement is shown for the base pretrained model vs. after offline RL (DRO) fine-tuning on each dataset.

| Dataset (Property) | Pretrained IgLM (no RL) | | | Offline DRO (IgLM; $\beta=0.01$) | | |
| --- | --- | --- | --- | --- | --- | --- |
| | $\rho \uparrow$ | P-value ↓ | IgLM LL ↑ | $\rho \uparrow$ | P-value ↓ | IgLM LL ↑ |
| MITLL AlphaSeq (Binding) | -0.09 | <0.001 | -3.99 | 0.45 | <0.001 | -5.59 |
| CR9114 Wisconsin (Binding) | -0.10 | <0.001 | -3.83 | 0.69 | <0.001 | -8.02 |
| CR9114 Caledonia (Binding) | -0.06 | <0.001 | -3.84 | 0.90 | <0.001 | -3.90 |
| Trastuzumab class (Binding class) | 0.17 | <0.001 | -4.41 | 0.44 | <0.001 | -3.53 |
| G6 expression (Expression) | -0.01 | 0.68 | -4.18 | 0.41 | <0.001 | -4.73 |
| G6 binding (Binding) | -0.03 | 0.44 | -4.18 | 0.23 | <0.001 | -3.87 |
| CovAbDab SARS-CoV1 (Binding class) | -0.10 | <0.05 | -4.50 | 0.28 | <0.001 | -3.59 |
| D44.1 (Binding) | -0.10 | <0.05 | -3.53 | 0.43 | <0.001 | -3.29 |
| CovAbDab SARS-CoV2 (Binding class) | -0.11 | 0.16 | -4.61 | 0.28 | <0.001 | -3.06 |
| Trastuzumab binding (Binding) | 0.11 | 0.31 | -4.35 | 0.37 | <0.001 | -5.68 |
| Marks 2021 (Immunogenicity) | 0.17 | 0.27 | -4.23 | 0.28 | <0.01 | -4.75 |

## 5. Conclusion

Generative models trained to maximize likelihood, such as IgLM, offer powerful tools for antibody sequence design, especially when infilling variable regions like CDR loops. In this work, we demonstrate how IgLM can be fine-tuned using RL to optimize for functional properties in both online and offline settings. In settings where proxy structural rewards are available, online RL via PPO with KL regularization successfully improves these properties while retaining sequence plausibility under the base model. When only experimental measurements are available, offline DRO fine-tuning greatly increases the Spearman correlation between model log-likelihoods and real-world property labels, consistently converting weak or negative baseline correlations into moderate-to-strong alignment without substantially compromising likelihood scores.

These results establish that the same IgLM backbone can be effectively fine-tuned via PPO in simulated, structural reward scenarios, and via DRO in data-driven, experimental regimes. PPO enables responsive structural optimization with minimal deviation from the pretrained space, while DRO allows leveraging static datasets to guide sequence generation toward measurable functional outcomes.

While online RL requires access to proxy rewards, which may be noisy or computationally expensive, and offline RL depends on existing datasets that may not span all desired properties or may introduce selection biases, our framework offers a flexible dual-path approach. It integrates in silico optimization and retrospective learning under a unified reinforcement learning paradigm.

Looking forward, we plan to explore integration of richer structural or physics-based rewards (such as those from AlphaFold or Rosetta), and tackle multi-objective optimization. Our complete code, datasets, and trained models are publically available to support reproducibility and to foster further innovation in RL-guided antibody engineering.

## Acknowledgements

We thank Livermore Computing at Lawrence Livermore National Laboratory (LLNL) for the computational resources that enabled this work. Funding was provided by the LLNL Laboratory Directed Research and Development project 24-ERD-022. This work was performed under the auspices of the U.S. Department of Energy by LLNL under contract DE-AC52-07NA27344. LLNL-CONF-2008752.

## Code and Data Availability

The complete code, both training and inference, along with datasets are available at https://github.com/LLNL/protein_tune_rl.

## A Appendix

### A.1 Illustration of the Optimal Policy in a Toy Example

In this section, we illustrate the theoretical optimal policy in a toy example. Assume a reference policy defined by a Gaussian distribution:

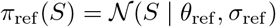

and a scalar reward function:

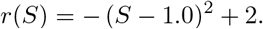

Given the objective

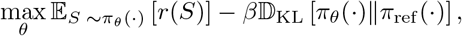

the optimal policy and soft value function are given by

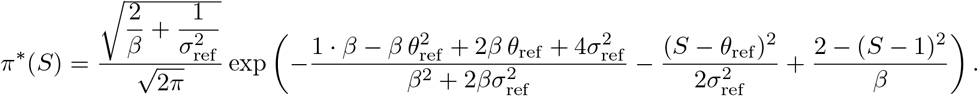

and

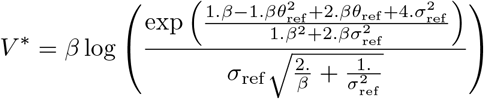

To visualize the impact of temperature *β*, Figure 4 plots *π*^∗^(*S*) as a function of *S* for several *β* values, with *θ*_ref_ = −0.7 and *σ*_ref_ = 0.1. As *β* decreases, *π*^∗^(*S*) concentrates more tightly around regions of high reward, demonstrating how temperature sharpens the optimal policy.

## APPENDIX A.2 Additional Online RL Results

### Density Visualizations for DRO Optimization

To better understand the behavior of the offline RL method (DRO), we visualize the distribution of generated sequences under two structure-based reward functions: beta sheet content and solvent-accessible surface area (SASA). Figure 5 shows histograms for both the target reward (top row) and the IgLM log-likelihood scores (bottom row) of sequences sampled from DRO-fine-tuned models.

**Figure 5.**
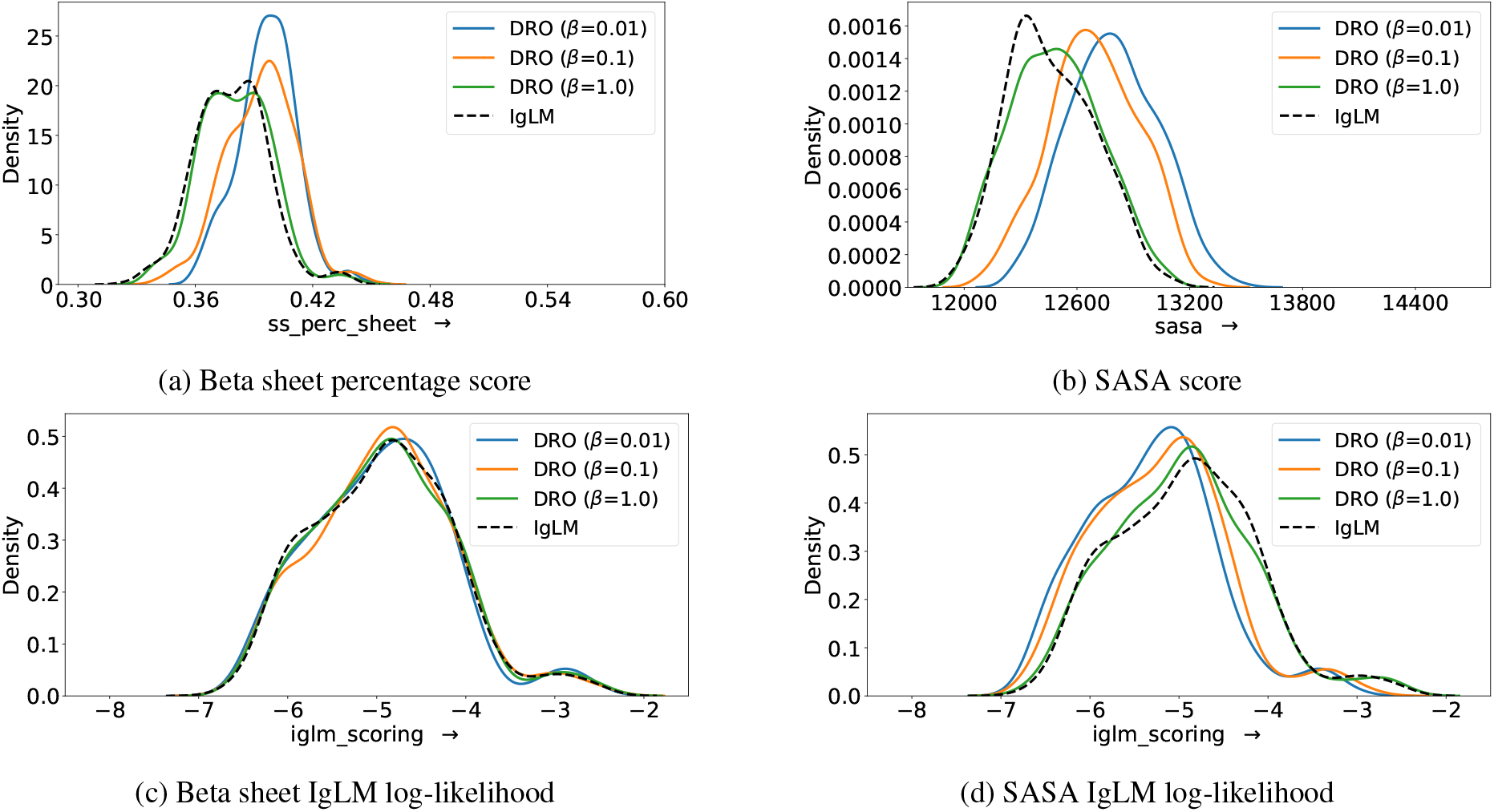
Distribution of reward scores (top) and IgLM log-likelihoods (bottom) on the evaluation set for models fine-tuned using DRO under structure-based rewards (Beta sheet and SASA).

For both objectives, DRO shifts the distribution of reward scores toward improved values relative to the original IgLM prior (not shown, but see Table 1 and 2). This confirms DRO’s ability to extract signal from offline reward data. However, the corresponding distributions of IgLM log-likelihoods remain centered around the prior, with only modest shifts. This reflects DRO’s balance between exploiting reward signals and preserving compatibility with the pretrained prior.

#### Pareto Analysis: Reward vs. Fluency Tradeoffs

To evaluate the balance between property optimization and language-model plausibility, we examine Pareto fronts plotting reward (e.g. percentage *β*-sheet or SASA) against IgLM log-likelihood. Figure 6 summarizes these trade-offs.

**Figure 6.**
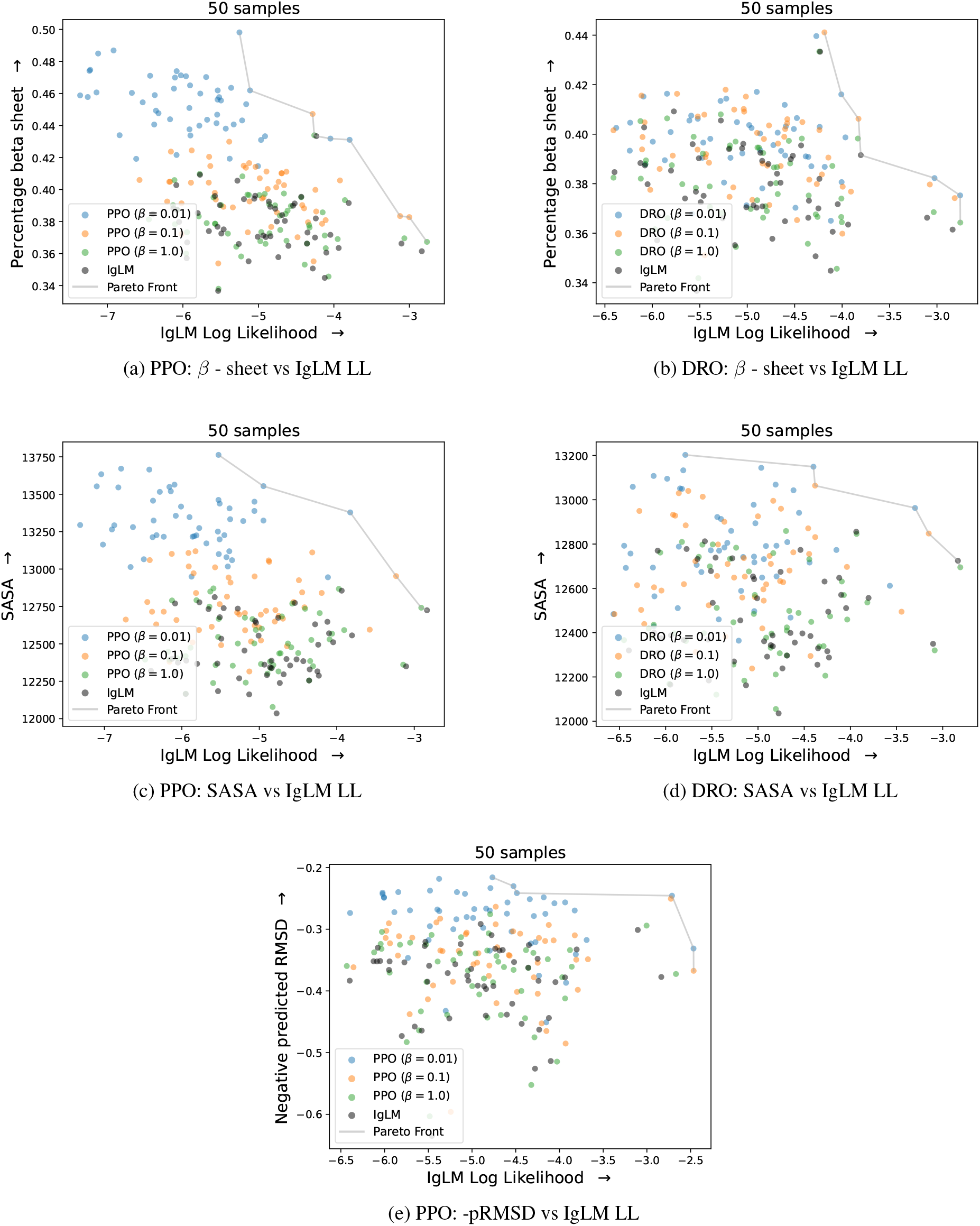
Pareto fronts showing reward versus IgLM log-likelihood across methods and tasks. Top row: PPO vs DRO for *β* - sheet content. Middle row: PPO vs DRO for SASA. Bottom row: PPO for pRMSD only. Each point represents the mean over 10 completions for the same prompt.

Each data point represents the average of 10 generated spans for the same context prompt. PPO’s behavior reflects an aggressive reward-centric strategy: stepping sharply off the pretrained distribution to gain functional improvement. DRO, by contrast, treads more cautiously—making reliable but conservative gains from static data. These observations hold consistently across tasks, confirming that PPO is ideal for strong online optimization, while DRO provides a safer, distributional-aware alternative when exploration is limited or prohibited.

## APPENDIX A.3 Offline RL with Real-World Antibody Datasets

In this experiment, we evaluate our offline RL framework using a diverse set of real-world antibody datasets-spanning binding-affinity assays, expression measurements, and binary classification labels (e.g. binders vs. non-binders), each drawn from distinct experimental studies [8]. Table 5 compiles all datasets, detailing their experimental origin, assay type, quantitative properties, and sample sizes.

**Table 5.** Overview of experimental datasets used in this study.

| Dataset | Property | Total | Description | Source |
| --- | --- | --- | --- | --- |
| MITLL AlphaSeq | Binding | 154632 | AlphaSeq assay scores approximating dissociation constants ($K_D$, nM, log scale); lower values indicate stronger binding. | [11] |
| CR9114 Wisconsin | Binding | 65535 | Binding affinities ($-\log K_D$) for CR9114 variants against Influenza A/Wisconsin/67/2005 (H3N2). | [46] |
| CR9114 Caledonia | Binding | 65094 | Binding affinities ($-\log K_D$) for CR9114 variants against Influenza A/New Caledonia/20/99 (H1N1). | [46] |
| Trastuzumab class | Binding (class) | 38839 | Classification of trastuzumab CDR H3 designs as binders/non-binders to HER2 based on affinity thresholds. | [34] |
| G6 expression | Expression | 4275 | Enrichment ratio indicating expression level for G6.31 antibody variants targeting VEGF. | [20, 8] |
| G6 binding | Binding | 4275 | Binding affinities ($K_D$, nM) for G6.31 antibody variants targeting VEGF. | [20, 8] |
| CovAbDab SARS-CoV1 | Binding (class) | 2912 | Classification labels for SARS-CoV-1 binding from CovAbDab antibody sequences. | [29] |
| D44.1 | Binding | 2048 | Binding affinities ($K_D$, nM) for D44.1 antibody mutants targeting lysozyme. | [41, 8] |
| CovAbDab SARS-CoV2 | Binding (class) | 844 | Classification labels for SARS-CoV-2 Beta variant binding from CovAbDab antibody sequences. | [29] |
| Trastuzumab binding | Binding | 423 | Binding affinities ($K_D$, nM) for zero-shot designed trastuzumab CDR H3 variants binding HER2. | [34, 8] |
| Marks 2021 | Immunogenicity | 217 | Clinical immunogenicity (ADA response percentage) for therapeutic antibodies. | [7, 8] |

